# Direct evidence for prediction signals in frontal cortex independent of prediction error

**DOI:** 10.1101/346213

**Authors:** Stefan Dürschmid, Christoph Reichert, Hermann Hinrichs, Hans-Jochen Heinze, Heidi E. Kirsch, Robert T. Knight, Leon Y. Deouell

## Abstract

Predictive coding (PC) has been suggested as one of the main mechanisms used by brains to interact with complex environments. PC theories posit top-down prediction signals, which are compared with actual outcomes, yielding in turn prediction-error signals, which are used, bottom-up, to modify the ensuing predictions. However, disentangling prediction from prediction-error signals has been challenging. Critically, while many studies found indirect evidence for predictive coding in the form of prediction-error signals, direct evidence for the prediction signal is mostly lacking. Here we provide clear evidence, obtained from intracranial cortical recordings in human surgical patients, that the human lateral prefrontal cortex generates prediction signals while anticipating an event. Patients listened to task-irrelevant sequences of repetitive tones including infrequent predictable or unpredictable pitch deviants. The amplitude of high frequency broadband (HFB) neural activity was decreased prior to the onset of expected relative to unexpected deviants in the frontal cortex only, and its amplitude was sensitive to the increasing likelihood of deviants following longer trains of standards in the unpredictable condition. Single trial HFB amplitudes predicted deviations and correlated with post-stimulus response to deviations. These results provide direct evidence for frontal cortex prediction signals independent of prediction-error signals.

## Introduction

Predictive coding theories postulate that the ability to detect unexpected environmental events results from a comparison of the actual state of our sensory world with predictions based on contextual knowledge of statistical regularities [1, 2]. Through this process the brain iteratively optimizes an internal model of the environment based on the sensory inputs [3], leading to improved interaction with the environment or to prediction error (PE) signals if predictions are violated [4]. In audition, passive listening to unexpected deviant sounds interrupting the context provided by a sequence of repeated standard stimuli results in robust differences between deviants and standards [5, 6, 7, 8, 9]. The classic auditory mismatch negativity (MMN) difference signal has been interpreted as a PE-signal although the mechanisms underlying the generation of these response differences are still debated. Two mechanisms differing with respect to the degree of memory involvement have been proposed [10]. The “neural adaptation” hypothesis argues that repeated presentation of stimuli leads to an increasingly attenuated and hence adapted response of feature-responsive neurons [11]: the more frequent the repetition the stronger the adaptation. Consequently, responses to the more frequent standards are more attenuated compared to rare deviants. By this account, response differences between frequent and infrequent stimuli do not reflect separate PE signals but simply originate from differential degrees of stimulus-specific adaptation (SSA), which has been demonstrated in auditory cortex of rats and cats [12, 13]. In contrast to the sensory habituation proposal, the “sensory memory” hypothesis posits that response differences reflect genuine PE-signals due to a higher-level comparison process that detects a deviation from a stored neural “memory” [14]. According to one version of the sensory memory hypothesis every stimulus is compared retrospectively to the extant ‘memory model’ with no active pre-stimulus anticipatory predictions involved. A third mechanism proposes that active prediction involves a prospective process reflected in pre-stimulus anticipatory activity [15]. In this model, under uncertainty, the simplest prediction would be occurrence of the more frequent stimulus, and violation of this prediction would generate a PE-signal when a deviant stimulus occurs. All the above mechanisms can be conceived as predictive, but only the last mechanism involves preparatory or proactive pre-stimulus activity using available predictive information. Most predictive coding schemes suggest separate prediction and prediction-error neurons or activity, but separating the two in practice proved challenging (for a comprehensive introduction and review see Heilbron and Chait, 2017). Whereas most evidence for predictive coding comes from the ultimate prediction-error signals, evidence of the prediction signal itself is harder to find. One line of evidence capitalized on stimulus omissions. In these paradigms, expected stimuli are omitted and responses time locked to the expected stimulus are taken as evidence of pure predictive signals. However, as reviewed by Heilbron and Chait, the results are somewhat mixed and they are subject to interpretational ambiguity.

An important assumption of predictive coding is that the prediction is formed prior to the event. Hence, recording predictive signals prior to the onset of the stimuli would be strong evidence for prospective, active predictions. Here, we utilized the high temporal and spectral resolution of direct cortical recordings from subdural ECoG electrodes to compare frontal and temporal prediction signals in five patients listening to trains of task-irrelevant auditory stimuli in two conditions. The conditions differed in the predictability of deviation from repetitive background stimuli. In *regular* sequences every deviant followed exactly 4 standards, whereas in *irregular* sequences deviants were randomly embedded in trains of standard stimuli. Subjects were instructed to ignore the sounds and watch a visual slide show. In a previous report, we concentrated on post-stimulus activity variations as a response to fully predictable and unpredictable deviants using the same data set [16]. Here, we focused on the pre-stimulus amplitude variations as metrics of the prediction of future stimuli and asked whether pre-stimulus activity signals regular deviations prior to their onset.

In the regular condition the occurrence of standards and deviants are fully predictable and we hypothesized that prediction of deviants will be different from the prediction of standard tones. In the irregular condition the occurrence of deviants cannot be predicted reliably. However, the increasing number of sequential standards may determine the likelihood of the occurrence of deviants: an imminent event (deviant) becomes more likely to occur the longer it hasn’t occurred. Reliance on this “hazard function” implies that standards and deviants and their overall probabilities are stored in memory. In contrast, in an opposite process the system expects ‘more of the same’ (inertia) and so with increasing number of sequential standards the expectancy of deviants would fade away. We hypothesized that the frontal cortex, assumed to be sensitive to higher order regularities would be differentially affected by the periodic vs. non-periodic occurrence of deviants. Specifically, we hypothesized frontal pre-stimulus activity would differ between pre-standard and pre-deviant windows in the regular condition, but not in the irregular condition. In the irregular condition frontal pre-stimulus activity would differ as a function of the number of standard tones presented, reflecting the hazard function. Following the results of our previous study, we expected the pre-stimulus activity in temporal cortex to be insensitive to stimulus regularity or to the hazard function.

## Methods

### Patients

Five epilepsy patients (mean age 33, SD = 9.23) undergoing pre-surgical monitoring with subdural electrodes participated in the experiment after providing their written informed consent. Experimental and clinical recordings were taken in parallel. Recordings took place at the University of California, San Francisco (UCSF) and were approved by the local ethics committees (“Committee for the Protection of Human Subjects at UC Berkeley”). The analysis of the post-stimulus effects from these patients with the same dataset was previously reported in [16].

### Stimuli

Participants listened to stimuli consisting of 180 ms long (10 ms rise and fall time) harmonic sounds with a fundamental frequency of 500 Hz or 550 Hz and the 3 first harmonics with descending amplitudes (−6, −9, −12 dB relative to the fundamental). The stimuli were generated using Cool Edit 2000 software (Syntrillium, USA). The stimuli were presented from loudspeakers positioned at the foot of the subject’s bed at a comfortable loudness.

### Procedure

While reclined in their hospital bed, participants watched an engaging slide show while sound trains were played in the background. Sound trains included high probability standards (p = 0.8; f_0_ = 500 Hz) mixed with low probability deviants (p = 0.2; f_0_ = 550 Hz) in blocks of 400 sounds, with a stimulus onset asynchrony (SOA) of 600 ms. In different blocks, the order of the sounds was either pseudorandom, with a minimum of three standard tones before a deviant (irregular condition), or regular, such that exactly every fifth sound was a deviant (Fig. 1A). Thus, in the regular condition, standards and deviants were fully predictable, whereas in the irregular condition, exact prediction was not possible. In both conditions the participants were instructed to ignore the sounds and watch a slide show of a variety of visual images changing in an unpredictable slow pace (∼ 3 sec per picture, unsynchronized with the auditory stimuli).

**Figure 1:**
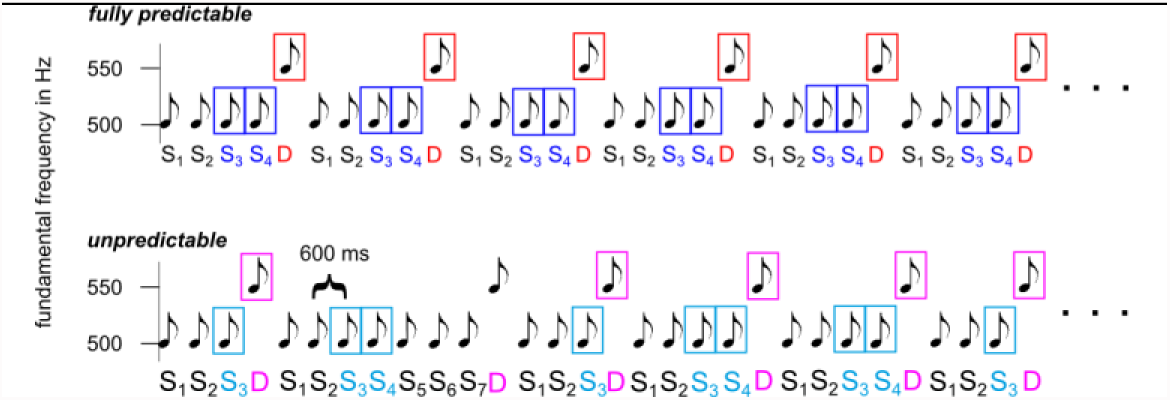
Paradigm. Participants watched a slide show while hearing passively sequences of sounds. High-probability standards mixed with low-probability Deviants were presented either unpredictably or were fully predictable ( exactly every fifth sound was a deviant). Standards (S_1-n_) are numbered based on their position relative to the previous deviant. Only standards following at least two standards were used for analysis (marked by rectangles)

### Data recording

The electrocorticogram (ECoG) was recorded at UCSF using 64 platinum-iridium-electrodes grids arranged in an 8 × 8 array with 10 mm center-to-center spacing (Ad-Tech Medical Instrument Corporation, Racine, Wisconsin; see **Figure 2** for grid location)). Grids were positioned based solely on clinical needs. Exposed electrode diameter was 2.3 mm. The data were recorded continuously throughout the task at a sampling rate of 2003 Hz.

**Figure 2:**
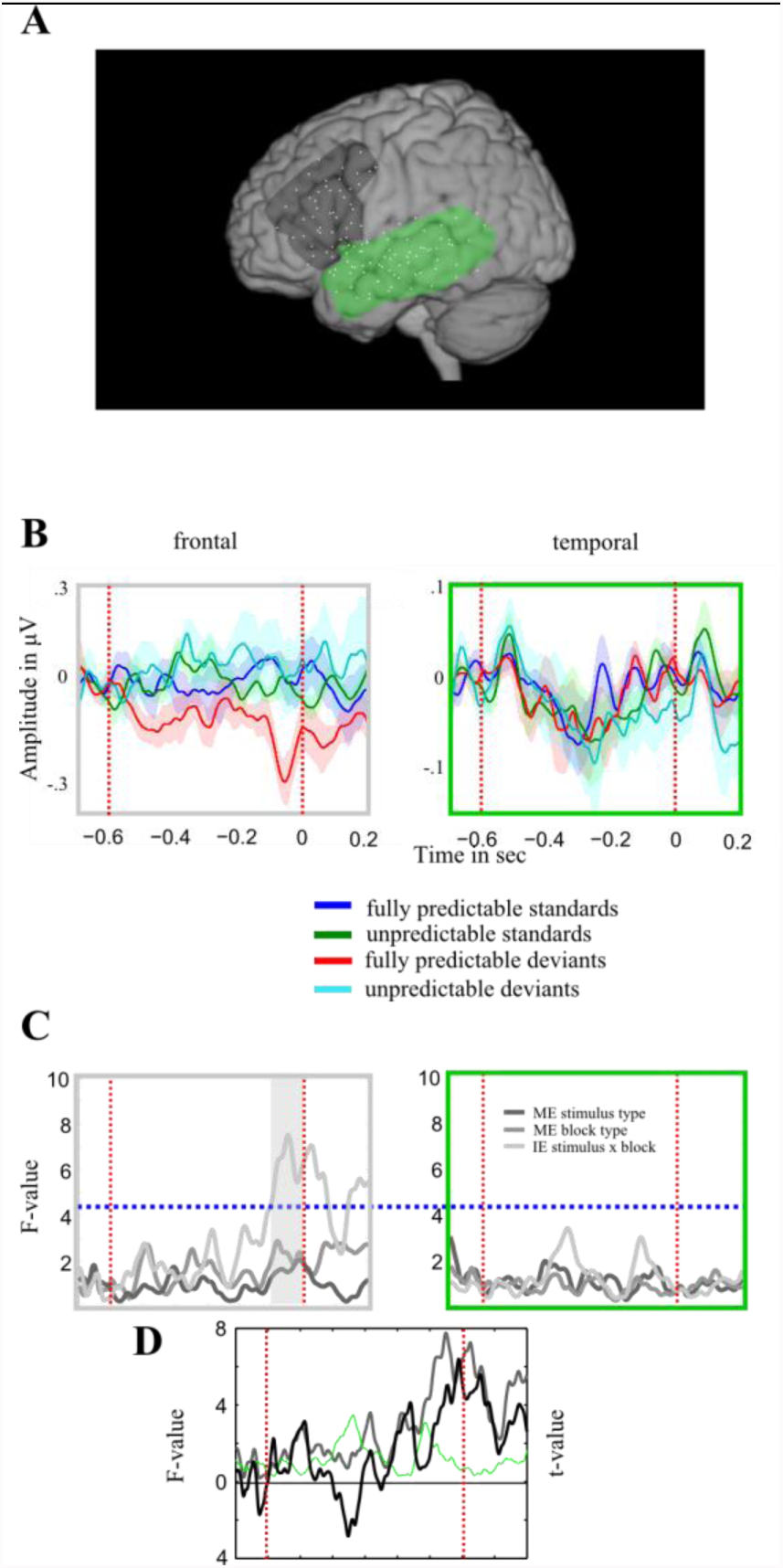
Time resolved analysis of variance. ***A*** Frontal (gray) and temporal (green) region of interest. ***B*** Baseline corrected HFB amplitude modulations prior to the standards and deviants in the regular and irregular condition. The shaded areas denote the standard error across channels. ***C*** Mean F (interaction/main effects) time series of channels loading highly on the frontal and temporal F_interaction_ 1st principal component, respectively. The horizontal dashed blue line indicates the critical F_interaction_ value based on permutation. The gray shaded area in the left panel of indicates the temporal interval of significant interaction. This starts ∼100 ms before stimulus onset lasting until the predictable deviant is presented. **D** shows both F_interaction_ time series (light gray lines in B right and green line in B left) together with the t-value (black line) showing the difference between the two ROIs in the degree of interaction between Type and Block. Frontal cortex shows stronger interaction before stimulus onset.

### Preprocessing

We used Matlab 2013b (Mathworks, Natick, USA) for all offline data processing. All filtering was done using zero phase-shift IIR filters. We excluded channels exhibiting ictal activity or excessive noise from further analysis. In the remaining “good” channels (see **Table** 1) we then excluded time intervals containing artifactual signal distortions such as signal steps of pulses by visual inspection. Finally, we re-referenced the remaining electrode time-series by subtracting the common average reference

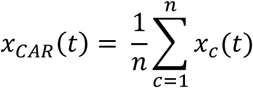

calculated over the *n* good channels *c* from each channel time series *x_c_*. The resulting time series were used to characterize brain dynamics over the time course of auditory stimulus prediction. For each trial (−1 sec to 2 sec around stimulus onset – sufficiently long to prevent any edge effects during filtering) we band-pass filtered each electrode’s time series in the HFB range (60-180 Hz; see *Supplementary Material*). We obtained the analytic amplitude *A_f_*(*t*) of the HFB frequency by Hilbert-transforming the filtered time series. We smoothed the analytic amplitude time series such that amplitude value at each time point *t* is the mean of 10 ms around each time point *t*. We then baseline-corrected by subtracting from each data point the mean activity of the −700 to −600 ms preceding the stimulus onset (i.e. 100 ms prior to trial N-1) in each trial and each channel. Channel time series were used for the following analysis steps that are explained in more detail below. We first parameterized the prediction of upcoming stimuli using an ANOVA (*I-Estimation of prediction*). In the next step we assessed the involvement of frontal or temporal cortices in this prediction effect (*II - Comparison between temporal and frontal cortices*). We used Receiver Operating Characteristics (ROC) analysis to test whether pre-stimulus amplitudes of channels predicted the upcoming stimulus on a single trial level (*III* – *Single trial ROC analysis*). Finally, we tested for an increasing predictability of deviants in the irregular condition following longer trains of standards (*IV – Increase of predictability as a function of train length*).

**Table 1.** classification results for ROC and SVM analysis and corresponding p values within the empirical distribution for cortical sites exceeding the significance threshold

|  | dorsal frontal | ventral frontal | dorsal temporal | ventral temporal |
| --- | --- | --- | --- | --- |
| **Channel level** |  |  |  |  |
| *fully predictable* |  |  |  |  |
| max AUC | **.63; p = .0003** | .56 | .53 | .56 |
| *irregular* |  |  |  |  |
| max AUC | .58 | .59 | .56 | .57 |
| **Group level** |  |  |  |  |
| *fully predictable* |  |  |  |  |
| max AUC | **.59; p = .01** | .56 | .55 | .55 |
| *irregular* |  |  |  |  |
| max AUC | .56 | .54 | .56 | .54 |
| **SVM** |  |  |  |  |
| *regular* |  |  |  |  |
| max Acc | **.62; p = .0015** | .56 | .53 | .55 |
| *irregular* |  |  |  |  |
|  | .52 | .55 | .57 | .54 |

#### I Estimation of prediction

We first identified frequencies showing a prediction effect (see Supplementary material). Given the fixed repetition of 4 standards followed by a deviant in the regular condition the occurrence of both standards and deviants should be predictable. We assumed that in areas with predictive activity, the activity ***P*** prior to (expected) deviants should be different from the brain activity prior to frequent standards:

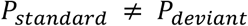

Conversely, since in the irregular condition the system does not know a-priori which stimulus will be heard the most frequent class (*standard tone*) is predicted, and, as a result, the activity ***P*** prior to the standards and deviants is equal:

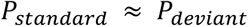

Statistically, the difference between conditions can be expressed as an effect of interaction using a 2-way ANOVA with the factors stimulus type (upcoming standard vs. upcoming deviant) and block type (regular vs. irregular), with the type effect being larger in the regular than irregular condition. We ran this 2-way ANOVA for each electrode (with trials as random variable), at every time point. This leads to 3 F-value time series (two main effects and one interaction: F_stimulus type_, F_block type_, F_interaction_) for each channel in the HFB band. The level of significance was corrected for multiple comparisons as described below.

Only deviants following the third and the fourth standard in a row (S_3_ and S_4_, respectively; see *Supplementary Material* for a full list of trials subjected to analysis) in the irregular condition were included in the analysis. All deviants following S_5_,…,S_N_ were excluded (see ***Figure 1*** and ***Supplementary Material***). This results in a pool of deviant trials which consist of regular deviants which always occurred after S_4_ and irregular deviants following S_3_ and S_4_. The pool of standard trials included only S_3_ and S_4_ trials in both the regular and irregular conditions. We did not include the first and second standards after a deviant, since during the pre-stimulus interval of S_1_ a deviant is presented and the pre-stimulus interval of S_2_ might still be influenced by the preceding deviant due to the short ISI. We excluded S_5_,…,S_N_ trials in the irregular condition since we hypothesized that the occurrence of deviants would be increasingly expected due to the hazard function. That is, we hypothesized that while longer trains of standards in the irregular condition increase the local probability of the standard, the occurrence of deviants also becomes more likely: since a deviant has not occurred for an extended sequence of events, its likelihood increases (“hazard function”). By not including irregular deviant following S_5_,…,S_N_ we also made the conditions more comparable for analysis, as in the regular conditions deviants never appeared after 5 or more standards. We focused on high frequency broadband (HFB) amplitude, which in our previous study showed earlier post-stimulus deviation signals than low-frequency ERPs [16] and differentiated between fully predictable and unpredictable deviation in frontal and temporal cortex (for representation of prediction signal in the whole time-frequency spectrum see *Supplementary Material*).

#### II Comparison between temporal and frontal cortices

We tested whether the F_interaction_ effect is localized to the temporal or the frontal cortex in the following way. The F_interaction_ time series were calculated in all channels separately over frontal and temporal regions of interest (ROI). Using a Principal Component Analysis (PCA) we estimated the typical course of F_interaction_-values across time separately for temporal and frontal channels. Channels loading highly on the first principal component are those that exhibit the strongest variation in terms of interaction amplitude across time. We chose the channels for which the Pearson correlation r with the principal component exceeded the 75^th^ percentile of all positive r-values. We set this level as a trade-off between a higher statistical power of a smaller number of channels and a stronger generalization across the cortex with a higher number of channels. We averaged the F_interaction_-values in these channels and checked whether the averaged F_interaction_-values in each region exceeded the empirically determined threshold derived from a surrogate distribution. This surrogate distribution of the interaction effect was constructed by randomly reassigning the labels (standard, deviant, regular, irregular) to the single trials in 1000 permutations for each channel. This leads to 1000 surrogate F_interaction_ time series. Significance criterion was a F_interaction_-value with p< .01 within the surrogate distribution of all F_interaction_ values. We next compared F_interaction_ effects between frontal and temporal electrodes with an unpaired t-test at each time point between the two groups of electrodes (frontal vs. temporal). To determine significance, in 1000 runs we randomly reassigned the labels (temporal vs. frontal) and applied the unpaired t-test.

#### III Single trial ROC analysis

We used Receiver Operating Characteristics (ROC) analysis to test whether pre-stimulus FFB amplitude predicted the upcoming stimulus on a single trial level [40]. We computed the predictive index that approximates the probability with which an ideal observer can predict the upcoming stimulus (standard sound vs. deviant sound) from the HFB amplitude on a single trial, separately in the regular and irregular condition. This index was estimated at each time point for each channel selected in step II as the **A**rea **U**nder the ROC **C**urve (AUC), based on the distributions of single trial amplitudes for deviants and standards, yielding an AUC time series for each channel. Specifically, in both the regular and the irregular conditions deviants were compared against the entire pool of standards. The AUC time series were subjected to PCA to find the course of AUC across time separately for four regions of interest of the lateral cortex (dorsal frontal, ventral frontal, dorsal temporal and ventral temporal cortex). In each region we determined channels loading highly on the respective first principal component and averaged the AUC time series across these channels, pooled across all subjects. We tested for significant deviations of the averaged AUC time series from predictive index at chance level (50%) using a permutation test. For this test, the empirical distribution of the main effect was constructed by randomly reassigning the labels (standard, deviant) to the single trials in 1000 permutations. We evaluated the statistical significance of the predictive index for each time point in each ROI in two ways. First, for each region and time point, to be considered as significant prediction, the averaged predictive index had to exceed the 95^th^ percentile of the empirical distribution. Second, to assess the difference between regions, we used a time point-by-time point 2 way ANOVA to test for differences of AUC values between 4 regions x 2 conditions (regular vs. irregular) across channels, followed by simple contrasts between regions in each condition. Again, we determined an empirical significance threshold for F values by randomly reassigning the four ROI labels in 1000 permutations of the same time point-by time point ANOVA.

The above analysis pooled across subjects to achieve a higher power at the expense of generalization. However, we also tested statistical significance on a group level with subjects as random variable. For that aim, in each subject, in each of the four regions and in each condition, we chose the channels with the maximal AUC value in the pre-stimulus interval (−600 to 0 msec) and compared their mean against a surrogate distribution.

Finally, to further assess the significance of our results we trained a support vector machine (SVM) on the same HFB amplitude time series separately for each of the four predefined ROIs in the regular condition. We estimated a prediction rate at each time point in each ROI determined by applying 10-fold cross-validation (CV). In each fold of the CV we randomly selected 10% of the dataset as test set and used the remaining 90% of trials for training a linear SVM. Since SVM is susceptible to an imbalance of class sizes (80% standard trials vs. 20% deviant trials), we balanced the number of data samples by randomly discarding standard trials. The feature space (HFB amplitude values) used for classification at each time point was constructed by randomly drawing observations (amplitude values) from all available observations (all amplitude values before standards and deviants) of each channel. SVM was then used to determine whether a given test sample was a pre-standard or pre-deviant stimulus trial. This procedure was repeated for each fold and results in one prediction for each of the data samples. For each time point the CV was repeated 100 times to estimate a distribution of prediction rates. To estimate an empirical significance threshold (confidence interval) under the null hypothesis of no predictive value, the same CV procedure was repeated 100 times again at each time point using randomly assigned class labels. Therefore, HFB activity in time intervals in which the observed prediction accuracy exceeded the empirically determined significance threshold distinguished between standard and deviant trials prior to their onset.

#### IV Increase of predictability as a function of train length

Throughout the experiment we pseudo-randomly varied the train length of standards in the irregular condition. We directly tested whether predictability varies as a function of train length in the irregular condition, congruent with a hazard function. We hypothesized that longer standard trains would result in stronger amplitude modulation of HFB before the occurrence of deviants. Specifically, we correlated the HFB preceding deviants with the length of the standard train before deviant in the irregular condition. While in the previous analysis we only used deviants following S3 and S4 here all deviants entered the analysis. To assess significance, Pearson’s correlation coefficient of each channel was compared against a surrogate distribution. This surrogate distribution was constructed by randomly reassigning the actual train lengths of single trial pre-deviant HFB values in 1000 runs. For each channel the confidence intervals (CI; 99.5%) of a normal distribution were determined.

## Results

### Comparison between temporal and frontal cortices

We studied 287 channels across all subjects, 120 were centered over frontal and temporal cortex. The HFB power was subject to a stimulus type (pre-deviant, pre-standard) x Block Type (regular, irregular) ANOVA at every time point and we evaluated the interaction term (F_interaction_) as a signature of predictive activity, separately for all frontal (N_frontal_ = 54) and all temporal channels (N_temporal_ = 66; see gray and green areas, respectively, in **Figure 2A**). Within each region we kept the channels loading highly on the first temporal principle component of the F_interaction_, time series, and compared their mean with the empirical surrogate distribution (Step I of data analysis in methods; **Figure 2B**). Frontal HFB (N_elec_ = 7) activity showed significant F_interaction_ values (maximal F_interaction_ = 7.76 p<.00001, at −51.4 ms) with neither a significant effect of stimulus type (maximal F_stimulus type_ = 2.44) nor of block type (maximal F_blocktype_ = 3.36) (left panel in **Figure 2B**). Temporal activity (N_elec_ = 10) did not show significant F values for any of the three effects (maximal F_stimulus type_ = 3.37; maximal F_block type_ = 2.02; maximal F_interaction_ = 3.47) (right panel in **Figure 2B**). The high F_interaction_ values in frontal cortex correspond in time with a decrease of HFB amplitude in the time range before the onset of deviants relative to the onset of standards in the regular blocks (where deviants and standards were predictable) but not in the irregular blocks (**Figure 2D**). The difference in F_interaction_ effects between frontal and temporal electrodes was significant (t_15_ = 6.49, permutation based p < .00001 at −11.5 msec; see **Figure 2C**).

### Single trial ROC analysis

The previous analysis informed us that the frontal cortex indexes, on the average, the upcoming onset of deviants in the regular and irregular condition differently by means of HFB amplitude modulation. We next analyzed whether pre-stimulus amplitudes predicted the upcoming regular deviant on a single trial level. To further localize neuroanatomical effects we subdivided the frontal and the temporal ROI into a dorsal frontal (around middle frontal gyrus), ventral frontal (around inferior frontal gyrus), dorsal temporal (around superior temporal gyrus) and ventral temporal region (around middle and inferior temporal gyri) (see **Figure 3A**). We computed the predictive index that approximates the probability with which an ideal observer can predict the stimulus (standard sound vs. deviant sound) from the HFB amplitude on a single trial level in both the fully predictable and irregular conditions. Only dorsal frontal channels exceeded the 99% significance threshold (AUC_ci99_ = .595) in the time range between −100 and −10 msec before onset of the regular deviant (see **Figure 3B** and **Table 1** for an overview of maximal AUC values for each region and for each condition). In contrast, we did not find a significant prediction of deviants in the irregular condition in any ROI following short trains of standards.

**Figure 3:**
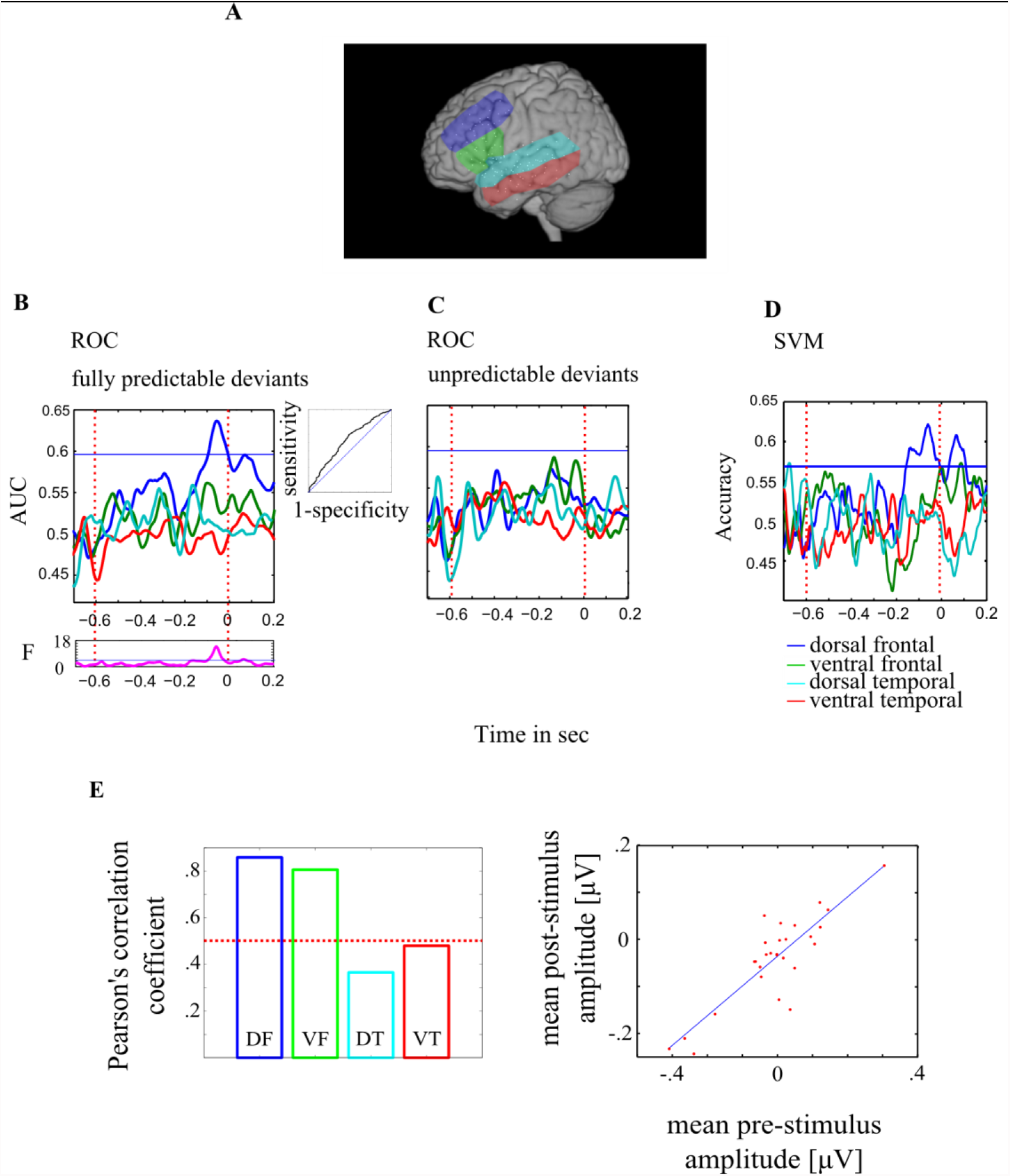
Pre-stimulus Hγ amplitude modulation predicts regular deviants on a single trial level in frontal cortex: **A** *Regions of interest*. The colors correspond to the colors in panels B-D. ***B-C*** Area under the curve (AUC) of the Receiver Operation Characteristics Curve (ROC) for predicting predictable (B) and unpredictable (C) deviants. Chance AUC is 0.5 and the horizontal line depicts the critical AUC for significance based on permuation analysis. The lower panel in B shows main effect of ROI in the regular condition ***D*** shows support vector machine classification results. ***E*** Correlation between prestimulus and poststimulus HFB amplitude across channels over the 4 ROIs. Left: Pearson’s correlation values for each of the four regions. The dashed line gives the 99% confidence interval of the surrogate distribution. Right: covariation of pre-stimulus and post-stimulus amplitude of electrodes over the more dorsal frontal ROI. Each dot represents one electrode. The blue line shows the linear fit to the data.

#### AUC differences between ROI

A time-point-by-time-point condition (regular vs. irregular) x ROIs (dorsal, ventral frontal, dorsal, ventral temporal) ANOVA with AUC as dependent variable revealed a highly significant difference of AUC values between ROIs (max F_ROI (3,19)_ = 10.4; p = 2.4^−11^ at −81 ms) and a significant difference of AUC values between conditions (max F_ROI_ (1,19) = 4.78; p = .0057 at −84 ms). Post hoc t-test (p = .05 corrected for 6 pairwise comparisons results; p_Bonferroni_ = .0083) revealed a significant difference between dorsal frontal AUC and dorsal temporal AUC (t_15_ = 4.42; p=.0005) and ventral frontal and dorsal temporal AUC values (t_14_ = 3.68; p=.002). Since our predictability model assumes differences in response before stimulus onset between the regular and irregular condition we carried out planned time point-by-time-point one-way ANOVA to test for differences of AUC between regions separately for the regular and irregular condition. This ANOVA revealed a significant difference between ROIs regarding the prediction of deviants in the regular condition (max F_ROI (3,10)_ = 17.8; p = .0002 at −59 ms), but not in the irregular condition (max F_ROI (3,10)_ = 3.8, n.s.). Post hoc pairwise comparisons in the regular condition revealed a significant difference between dorsal frontal and dorsal temporal (t_7_ = 5.42; p = .001), between dorsal frontal and ventral temporal (t_7_ = 8.1; p = .0012), between dorsal and ventral frontal (t_6_ = 4.8; p = .01) and between ventral frontal and ventral temporal AUC (t_7_ = 6.1;p = .009) with the strongest predictive value in the dorsal frontal region.

#### Group level

At the group level (i.e. with subjects as the random variable) we also found with planned comparisons the dorsal frontal cortex HFB activity predictive for the occurrence of deviants in the regular condition. AUC values exceeded the 95% ci (AUC_ci95_ **=** .57) in the dorsal frontal cortex (max AUC_mf_ = .59; p = .01 at −76 ms) in the time range between −95 and −57 ms before onset of the regular deviant but not in the irregular condition (see **Table 1**).

#### SVM

To support our results we also employed a SVM classifier. Accuracy exceeding the empirically defined significance threshold between −152ms and - 6ms (max Acc_mf_ = 62.5%; p=.0015 at −57.7 ms) was observed only for the predictable deviants, exclusively in the dorsal prefrontal cortex

In previous work (Dürschmid et al. [16]) we found that post-deviant HFB amplitude was reduced in the regular condition compared to the irregular condition. Since we now found that predictable deviants in the regular condition are heralded by a pre-stimulus HFB decrease, we tested if the two phenomena are correlated. That is, we tested whether a pre-stimulus HFB decrease corresponds with smaller HFB amplitude increase following deviant onset both across channels and across trials. First, in each ROI we correlated HFB amplitude preceding stimulus onset (average across −100 to 0 ms) with the amplitude following stimulus onset (average across 0 to 300 ms) across channels. The four resulting Pearson’s correlation values were tested against a surrogate distribution. This surrogate distribution was constructed by shuffling the order of channels in 1000 iterations. Hence, in each iteration we randomly yoked the pre-stimulus values with post-stimulus values and hence randomly reassigned the pre-stimulus value of one channel to post-stimulus value of another channel. The critical r-value denoting statistical significance was .5. Pre-stimulus amplitude correlated with post-stimulus amplitude in dorsal frontal cortex (r=.86; p = .000003) and in ventral frontal cortex (r = .8; p = .00001) but not in the temporal cortex (r<.5; see ***Figure 3E***). Next, we tested whether the pre-stimulus/post-stimulus relation is also true at a single trial level. Hence, we correlated within each electrode the average amplitude in the pre-stimulus and post-stimulus interval across trials. Each individual Pearson’s correlation coefficient was compared against a surrogate distribution and excluded if smaller than the critical value (*rcrit* =.1). This surrogate distribution was constructed by shuffling the order of trials in 1000 runs. Hence, in each run we randomly reassigned the pre-stimulus value of one trial to post-stimulus value of another trial by randomly permuting the pre-stimulus values. On average electrodes in the dorsal frontal cortex showed highest *r* values (MF: .39; IF: .38; ST: .27; MT: .30). Differences of pre/post correlation coefficients between ROIs were evaluated with an ANOVA. We found a significant main effect of ROI (F_(3,103)_ = 5.41; p=.0017). Post-hoc tests revealed a significant difference between dorsal frontal and dorsal temporal cortex (t_54_ = 3.35; p=.0015) and between ventral frontal and dorsal temporal (t_52_ = 3.27; p=.0019) at a Bonferroni corrected level (p_Bonferroni_ = .0083), and a trend in the same direction between dorsal frontal and ventral temporal cortex (t_48_ = 2.09; p=.04, uncorrected) between ventral frontal and ventral temporal cortex (t_46_ = 1.99; p=.05, uncorrected) with the strongest correlation in the both frontal regions – which could not be distinguished from each other – compared to temporal regions.

### Increase of predictability as a function of train length

The train length of standards in the irregular condition varied pseudo-randomly allowing us to test whether pre-stimulus predictive activity varies gradually as a function of train length. We surmised that two effects could be at play. Temporally local effects suggest that the probability of a standard tone increases the more standard tones are played in a row as the recent history becomes more stable. In contrast, using a more global strategy, the so-called “hazard function” suggests that, given that deviations will happen eventually, expectation of a deviant increases the longer it is since the last deviation. To test whether and where such effects prevail, we correlated pre-deviant HFB amplitude with train length of standards before deviants. The average correlation coefficient across all significant channels was -.23 (SD = .026). However, ***Figure 4*** shows that the direction of correlation between HFB amplitude and standard train length was different between temporal electrodes, showing positive correlations, and frontal electrodes, showing negative correlations. Individually, only the negative correlation in frontal channels reached the permutation critical r values of r_crit_ = ± .19,(white dots in ***Figure 4***). Considering that the analysis of the regular vs. irregular condition indicated that a decrease in HFB amplitude indicates proactive prediction of a deviant, these results suggest that frontal electrodes ‘apply’ predictions in the irregular condition based on the more global hazard function strategy.

**Figure 4:**
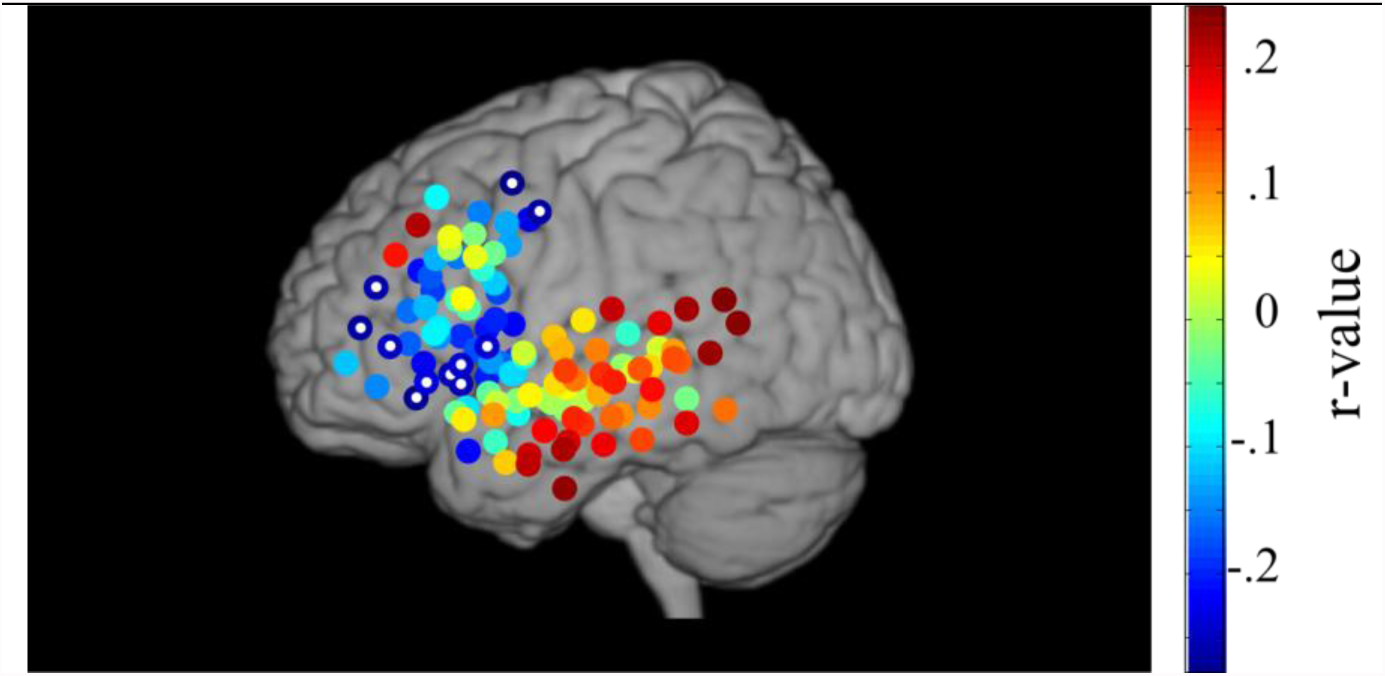
Prefrontal electrodes reflect the hazard function in irregular sequences. Each circle depicts channel positions with the color coding the Pearson correlation’s correlation coefficient between train length and pre-deviant HFB amplitude. Channels with a white dot show a statistically significant correlation. HFB amplitude significantly decreased after longer trains of standards in frontal cortex, while HFB amplitude tended to increase with longer trains of standards in temporal cortex.

## Discussion

Unexpected deviant sounds in a sequence of repeated standard sounds result in differential responses between deviants and standards. These mismatch signals may result from violations of prospective prediction of the next stimulus, formed on a moment-by-moment basis, or as a result of a retrospective comparator mechanism between a passively evolved statistical model of the context and a new input. We have previously shown that lateral frontal sites show reduced HFB response to predictable compared to unpredictable stimuli, whereas temporal sites are not sensitive to the deviant predictability [16]. Here, we asked whether predictable deviants are indexed differently than unpredictable deviants in terms of proactive, pre-stimulus activity modulation suggesting actual anticipation of the upcoming deviant. Finding such activity would provide evidence for predictive activity not confounded with stimulus-evoked prediction-error signals.

To address this proposal, we examined the role of lateral frontal and temporal cortices in generation of event-related prediction signals of regular (fully predictable) and irregular (pseudo-random) auditory events. Regular deviations were preceded by decrease in the power of the HFB in the frontal cortex but not over the temporal cortex. This power modulation was robust on a single trial level in the frontal cortex. Pre-deviant HFB modulation correlated with post-deviant HFB modulation across both channels and trials in the frontal cortex, indicating a possible causal effect of pre-stimulus HFB decrease on reduced response to predictable deviants (better prediction leading to less prediction-error). Finally, we found a strong inverse correlation between the train length of standards and the pre-deviant HFB decrease over frontal cortex, consistent with sensitivity to a hazard function, and a weaker direct correlation over the temporal cortex, consistent with sensitivity to local rather than global statistics in this region. Taken together, these results provide evidence for proactive, anticipatory processes in frontal cortex, based on global statistics, which may provide the basis for the curtailment of orienting response to predictable events in an unattended stream.

Predictive coding theory assumes that the brain uses available information continuously to predict forthcoming events and reduce sensory uncertainty [17]]. Recent studies gathered convincing evidence that stimulus-related responses depend on predictability of upcoming stimuli [18, 19, 20, 21, 22, 23, 24, 25, 26]. Fogelson et al [22] for example found shorter P3b latencies for predictable targets compared to non-predictable targets, which could be due to involvement of preparatory mechanisms shortening the duration of the stimulus evaluation. They further linked this mechanism to the prefrontal cortex as they found PFC lesioned patients to be impaired in the use of a short predictive series of visual stimuli to detect targets. Comparably, in Dürschmid et al [16] we showed that frontal HFB, but not temporal HFB, nor low frequency ERPs in either region, discriminated between predictable and unpredictable changes. This was indicated by a strong HFB mismatch responses (MMR) to irregular deviants but nearly no MMR for predictable deviations in frontal electrodes. That is, in frontal electrodes the HFB responses to predictable deviants were ‘quenched’.

Common to all these studies is the fact that prediction is studied indirectly, as the influence of predictability on the actual response to an event, presumably measuring the prediction-error signal or a mixture of prediction and prediction-error signals. In contrast, anticipatory effects of predictive coding are less explored. A step forward in dissociating prediction signals from prediction-error is provided by studies investigating the omission of stimuli in a sequence. Generation of a signal time-locked to an absent stimulus is taken as evidence for top-down, predictive coding, although whether they present predictions signals per se or prediction-error signals is debated (see Heilborn and Chait, 2017 for an up-to-date review and discussion). Most omission-locked responses can be considered as resulting from violation of a general prediction for the occurrence of a stimulus at a given time (a temporal prediction). Two studies directly examined whether predictions are more specific for the occurrence of a specific event. SanMiguel et al. [23] had subjects generate environmental sounds by pressing a button, with occasional button presses not producing sounds (omissions). Omission responses were found only when the same sound was repeatedly elicited by the button presses (and was thus predictable) but not otherwise. This result suggests specific predictions; however it was elicited in an active task and so does not speak for automatic, non-intentional predictive coding. In a passive task with visual distraction, [26] presented sequential pairs with a very short interval between them. The intra-pair frequencies were identical, whereas the frequencies roved between pairs. The results showed evidence for omission-locked responses when the identity of the omitted stimulus could be predicted (because it was the second sound in the pair), but not when only its timing could be predicted (because it was the first in the pair). However, subjects may have perceived each pair as an auditory object, and the omission of the second sound in the pair, which elicited the critical omission response, might be a response to a duration change.

Rather than looking at post-stimulus or post-omission responses, our current results addressed the pre-stimulus time, a time window at which no error may yet be computed, and thus the modulation of activity has to be ascribed to prediction per se. Similarly, Kok et al [27] decoded from MEG recordings the orientation of visual grating stimuli which could be predicted by a preceding auditory stimulus (valid visual stimulus) or not (invalid visual stimulus). Subtracting the orienting signal of valid from invalid gratings revealed differences before stimulus presentation suggesting actual proactive processing, in the form of pre-activating an anticipated sensory template. However, while the decoding approach used by Kok and colleagues using extra-cranial MEG data suggests the presence of information regarding the identity of an upcoming stimulus, it is short of localizing this predictive activity, nor delineating its nature. Our findings show that predictable deviants are preceded by HFB amplitude, and that this decrease is expressed in frontal cortex but not over the temporal cortex, corroborating a hierarchy of prediction in the human brain [16]. This hierarchy in line with the notion that whereas in early stages of processing information is represented based on bottom-up signals, in higher levels of cortical processing mainly deviations from expectation are registered while predictable components are filtered out [2].

Several studies have used dynamic causal modeling (DCM) of EEG or MEG to test the interaction between primary auditory sources, lateral superior temporal gyrus cortex, and inferior frontal cortex (IFG) in the generation of the mismatch negativity [28]. These studies suggested a hierarchical feedforward-feedback cascade in which the IFG sits at the top, providing top-down predictions to (and receiving PE signals from) the STG, which in turn provides top-down predictions to (and receives PE signals from) the early auditory cortex. Recently, Phillips et al., [29, 30] extended these models to multiple types of deviations measured concurrently, and validated the models, originally tested on MEG data, with ECoG data from two patients, one with electrodes on the left and one on the right hemisphere. Relevant to the current study, their studies added “internally generated prediction” signals to the predictive model, affecting the IFG, mostly on the left. As the models were based on the post-deviant activation, the latter signals may be related to the reduction of HFB in response to predictable deviants which we previously reported [16]. However, Phillips et al.’s models suggested that the prediction signal affecting the IFG is limited to temporal deviations (duration deviations and gaps in their study), but not pitch, intensity or location deviations, whereas our findings showed clear effects of predictability when the deviation was in pitch. Moreover, as noted, the predictive signal suggested by the models was based on the post-deviant activity, during the elicitation of MMN, whereas the current study shows pre-stimulus, anticipatory modulations of activity in the frontal cortex, correlated with the post-deviant effects. Future models will need to incorporate these anticipatory effects.

Indeed, one of the novel observations in this study is the existence of pre-stimulus predictive effects which do not entail temporal predictions per se. That is, the suppression of HFB power indexed the identity (standard or deviant) of the predicted stimulus rather than its specific timing. Moreover, this was observed in a paradigm in which all stimuli were task-irrelevant, did not require a response, and did not carry any reward value. Previous findings of anticipatory response typically involved active preparation for an upcoming imperative stimulus, reflected in the contingent negative variation (CNV) recorded on the scalp [31, 32], or in reward-prediction signals of different types [33]. The current finding provides evidence for ongoing, task-independent, anticipatory predictive signals, operative even before the stimulus was received. The correlation we found between the pre-stimulus HFB suppression and the post-stimulus deviant-related response suggests that the pre-stimulus activity may shape the initial response to the incoming stimulus although the causality remains to be proved.

What is the relationship between our findings of predictive, pre-stimulus modulation of the HFB signals, with the accounts of the mismatch response elicited by the deviant? Post-stimulus effects like the MMN may involve different states of neural adaptation. Thus, with repeated presentation of stimuli with a given feature value (e.g. pitch or location), neurons tuned to this feature value will get adapted. This creates a model of the recent history, and under an assumption of stationarity, provides a reasonable prediction of the future environment [11]. Other models [14] suggest that beyond adaptation, the repetition of a stimulus increases the absolute excitability of neurons tuned to values not included in the repeated stimulus. By both accounts, when a new stimulus arrives, it will elicit a stronger response, sometimes described as a comparison of the new stimulus with the current model which generates a prediction-error signal. The current findings of predictive pre-deviant modulation of activity cannot be explained by either mechanism. First, we compared the response to deviants following a similar number of standards in the random and predictable conditions, and overall deviants and standards had the same probability in both conditions. Thus, either adaptation or lateral excitation should have been similar across conditions. Second, since the effect occurred before the deviant, it cannot be due to activation of non-adpated/excited neurons sensitive to the pitch of the deviant or by a process of comparison. Instead, it seems to represent an actual high level prediction, modifying the post-stimulus comparison between the actual input and the ongoing prediction, which is based on the local probabilities.

Our main experimental manipulation, the comparison between completely predictable and randomly placed deviants, pointed to HFB reduction as a signature for predicting a deviation. This observation allowed us to investigate, post-hoc, whether anticipatory predictions are generated during irregular, random sequences as well. We found that in frontal cortex, pre-stimulus HFB decreased as the train of deviants became longer. Under the premise, derived from the results of the main comparison between regular and irregular conditions, that HFB reduction reflects increasing likelihood of a deviant, this pattern matches well to the so-called “hazard function”, in which an imminent event becomes more likely to occur the longer it hasn’t occurred. This suggests that the frontal cortex predictive capacity is not limited to highly structured sequences like our regular sequence, but rather, that frontal cortex generates complex predictions based on global probabilities, even in a task-irrelevant irregular stream. This progressive increase in deviant prediction seems akin to the progressive increase in the contingent negative variation (CNV) as a function of distance from the last deviant reported by [34]. Note however that the CNV effect was seen in that study only when subjects were required to attend the stimuli and especially to the deviants, whereas in our case stimuli were task irrelevant. The temporal cortex electrodes in our patients elicited a trend towards an opposite effect compared to the frontal ones – pre-stimulus HFB activity increased its amplitude the longer the standard train was. This is consistent with the notion that temporal cortex is based on recent history, such that with longer standard trains, “more of the same” is expected.

The current findings are consistent with our previous report, showing that predictions based on the global statistics of the sequence are stronger and possibly restricted to the frontal recording sites, whereas the temporal recording sites reflect only the local effects. Under the predictive model paradigm, PE signals should be carried forward to higher nodes in the network, to allow modification of the current model and influence the next prediction. However, taken together, our previous and current findings highlight the complex nature of this process, which has to entertain multiple levels of possibly conflicting predictions. Just prior to a deviant in the regular condition, and also after a long train of standards in the irregular condition, processes based on local effects predict another standard, whereas predictions based on the global statistics predict a deviant. In this situation, it seems efficient to prevent PE signals elicited at the temporal (auditory) cortex from propagating up the hierarchy and modifying a veridical model of the environment. Similarly, it seems that the prediction of an upcoming deviant based on global statistics, present at the frontal cortex, does not propagate down the network to mitigate the prediction-error signal invoked by the expected deviant in the temporal cortex [35]. Possibly, detecting local changes would be advantageous for parsing auditory input into meaningful chunks, but would require too many resources if every change would lead to a behavioral response such as a shift of attention. Our results suggest that the flow of information up and down the hierarchy of the network is not as simple as may be gleaned from typical DCM diagrams.

Predictable deviants were heralded by a robust HFB decrease, which correlated with reduced stimulus evoked responses in frontal cortex. Previous studies have shown pre-stimulus modulation of neural activity during selective attention tasks. These changes usually involve activation (increased firing rate in animals, increased BOLD response) prior to task relevant stimuli [36] [37] and deactivation prior to task irrelevant stimuli [38] [39]. Our study did not use a classic selective attention task, but involved a competition between the primary task the subjects were engaged with (viewing a slide show) and the potential distraction caused by the auditory stream, especially by deviant events. Thus, the HFB decrease observed prior to an expected deviant could reflect the same filtering mechanism previously observed during selective attention. Under this premise, the current results suggest that this inhibitory anticipation can be generated selectively, and in predictive manner, in an unattended stream.

Taken together, the pre-stimulus and post-stimulus HFB response reveals a unique role for prefrontal cortex in utilizing global regularity to control responses to deviant stimuli. Frontal HFB selectively signals upcoming regular deviants with a decreased amplitude prior to deviant onset. Subsequently, only unpredictable deviants elicit a strong HFB response, related to triggering an orienting response to an environmental perturbation. Our results highlight a selective role of frontal structures in actively computing predictions.

## Supplementary Material

To reveal predictive signals prior to the onset of the stimulus we evaluated high γ amplitude modulation in the prestimulus interval. The trials were grouped according to their identity in the poststimulus interval (see *Supplementary Table* 1 column 4). Column “Stimulus Type” and “Condition” give the group labels subjected to the ANOVA for the stimulus type (deviant vs. standard) and condition (regular vs. irregular), respectively. The 2 way ANOVA was conducted at each time point in the prestimulus interval (-.6 to 0 sec). In this prestimulus interval the last stimulus presented was always a standard (S_2_, S_3_, or S_4_). The high γ time series in the prestimulus interval were baseline corrected by subtracting the 100 msec preceding the prestimulus interval (last 100 msec following S_1_, S_2_ or S_3_ respectively). In this baseline interval we expect the prediction to be for a standard. Note that in the irregular condition deviants can occur following a train of only 3 deviants (see row **IV** of *Supplementary Table* 1). Hence, in the case shown in **V** of *Supplementary Table* 1 theoretically a deviant could be assumed in the baseline interval of the prestimulus interval, even though S_4_ actually follows. However, under the premise that all train lengths are evenly distributed the likelihood of a deviant following S_3_ is limited. Note also that the response to the stimulus presented prior to the prestimulus interval (stimulus onset at −600 msec) should be less influenced by repetition suppression as we limited the analysis to the deviants following S_3_ and S_4_ and excluded deviants following S_5_ and S_N_.

**Supplementary Table 1:** lists all trials subjected to the 2 way ANOVA.

|  |  |  | Prestimulus<br>interval<br>(ANOVA) | Poststimulus<br>interval | Stimulus<br>type | condition |
| --- | --- | --- | --- | --- | --- | --- |

| Temporal interval in sec |  | -1.2 to -.6 | -.6 to 0 | 0 to .6 |  |  |
| --- | --- | --- | --- | --- | --- | --- |
| Regular condition | I | S<sub>3</sub> | S<sub>4</sub> | D<sub>regular</sub> | 1 | 1 |
|  | II | S<sub>1</sub> | S<sub>2</sub> | S<sub>3regular</sub> | 0 |  |
|  | III | S<sub>2</sub> | S<sub>3</sub> | S<sub>4regular</sub> |  |  |
| Irregular condition | IV | S<sub>2</sub> | S<sub>3</sub> | D<sub>3irregular</sub> | 1 | 0 |
|  | V | S<sub>3</sub> | S<sub>4</sub> | D<sub>4irregular</sub> |  |  |
|  | VI | S<sub>1</sub> | S<sub>2</sub> | S<sub>3irregular</sub> | 0 |  |
|  | VII | S<sub>2</sub> | S<sub>3</sub> | S<sub>4irregular</sub> |  |  |
|  |  | S<sub>4...N-1</sub> | S<sub>5...N</sub> | D<sub>5...N</sub> | 1 |  |
|  |  | S<sub>3...N-2</sub> | S<sub>4...N-1</sub> | S<sub>5...N</sub> | 0 |  |

## Specificity of predictive code in the High frequency activity

In this study we focus on high γ activity since high γ activity signaled prediction errors earlier and distinguished between fully and unpredictable deviants. However, we additionally verified that a prediction signal operationalized as the F_interaction_ value is represented mainly in the high γ range in the following way. For each trial (−1 sec to 2 sec around stimulus onset – sufficiently long to prevent any edge effects during filtering) we band-pass filtered each electrode’s time series at 42 frequency bands (log-spaced between 3 and 200 Hz) with a bandwidth of 10% of the center frequency. We obtained the analytic amplitude *A_f_*(*t*) of each frequency *f* by Hilbert-transforming the filtered time series. We smoothed the time series such that the amplitude value at each time point *n* is the mean of 10 msec around each time point *n*. We then baseline corrected the prestimulus trial (N-1) activity by subtracting the mean activity from the −700 to −600 msec preceding the stimulus onset in each trial of each channel (100 msec prior trial N-1, as stated above this could be the last 100ms of S_1_, S_2_ or S_3_). This prediction signal can be equally high in different frequency bands but can be distributed across networks of different spatial extension and hence different sets of electrodes. Averaging across the whole set of all electrodes would favor frequency bands with a larger set of electrodes showing a high F_interaction_ value. Hence, in each frequency we averaged the F_interaction_-value across the prestimulus interval separately for each electrode (in the first step we used the −400 to 0 msec as the prestimulus interval to cut out the stimulus response but systematically varied this interval – see below). We then took the 5% of electrodes with the highest averaged F_interaction_-value in this prestimulus interval and averaged the F_interaction_-value time series across the selected channels. This results in one F_interaction_-value time series for each frequency which were tested for significance against a surrogate distribution. This surrogate distribution of the interaction effect was constructed by randomly reassigning the labels (standard, deviant, regular, irregular) to the single trials in 1000 permutations for each channel. This leads to 1000 surrogate F_interaction_–value time series for each frequency. Significance criterion was a F_interaction_-value with p< .01 within the surrogate distribution. In these channels showing the highest frequency specific F_interaction_ value we also compared F_stimulus-type_ and F_block-type_ time series across frequencies.

In the initial step of this analysis we chose a prestimulus interval ranging from −400 to 0 msec. The rationale was to separate any prediction signal (expected rather in the end of the prestimulus interval) from differences in response to the stimulus presented in the prestimulus interval. However, a shorter interval disadvantages low frequencies if one takes into account that at least 3 cycles of the underlying oscillation are necessary to evaluate an amplitude modulation. For example, the θ activity has a center frequency of 6 Hz and hence a cycle length of 166 msec. To fully cover 3 cycles an interval of at least 498 msec is necessary to evaluate amplitude modulation. In contrast, longer intervals decrease the possibility to detected transient fluctuations in higher frequencies. Therefore, we systematically varied the interval upon which we selected the 5% of best electrodes. We averaged F_interaction_ values in 5 different intervals (−600/−500/−400/−300/−200 to 0 msec) and assessed which frequency carries the highest predictive strength in terms of F_interaction_ value.

We found a significant effect of interaction in the Hγ range (Hγ: 60-180; see **Supplementary Figure 1A**). In all subjects we found channels exceeding the 95% confidence interval (*S1*: 9 channels; *S2*: 5 channels; *S3*: 3 channels; *S4*: 7 channels; *S5*: 5 channels). The empirical F_interaction_ value derived from the permutation was 4.28. We found highest F_interaction_ values in the time range between −400 and −300 msec (*F* = 5.201; *p* = .0015; *df =* 1,458) and between −100 and 0 msec (F = 5.29; p = .0012). Neither the averaged F_stimulus type_- nor the F_block type_ – time series of these channels reached the empirical significance threshold (see **Supplementary Figure 1 B**).

**Supplementary Figure 1:**
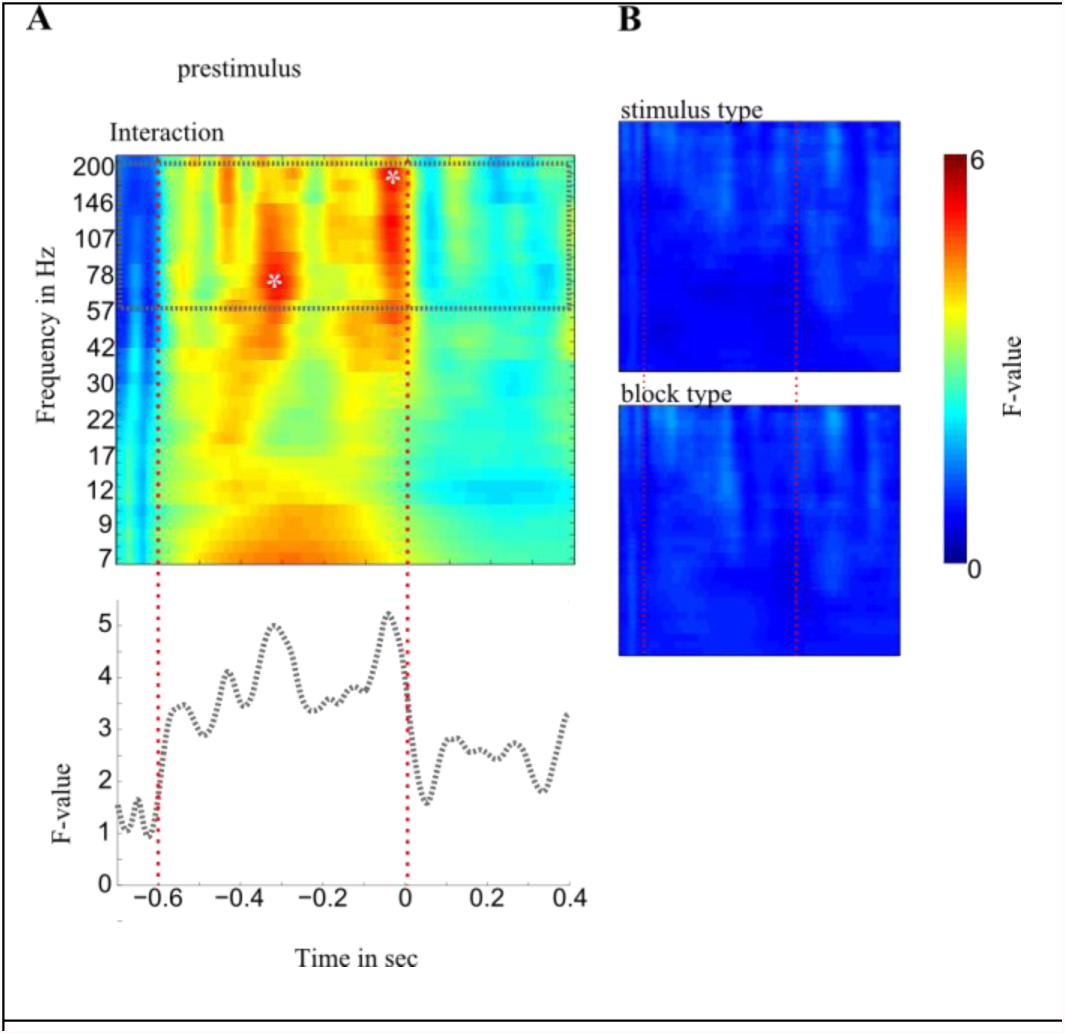
Depiction of selection of frequencies reflecting the onset of regular deviants. ***A:*** A time point-by-time point ANOVA compared prestimulus activity prior to stimulus and condition types (standards vs. deviants, regular vs. irregular) in each channel and each frequency band. In each frequency we averaged the F_interaction_-value across the prestimulus interval (−400 to 0 msec) separately for each electrode. F_interaction_ time series of electrodes with the highest averaged F_interaction_-value (>95% percentile of the distribution) were pooled together. Warm colors indicate high F-values and cold colors show small F-values. Only the F-values in the high gamma range exceeded the empirical significance threshold. The dashed gray line shows the averaged F_interaction_ time series across the narrow frequency bands spanning the high gamma range. ***B*** shows the F_stimulus type_ and F_block type_ time series averaged across channels with the highest F_interaction_ value shown in ***A***.

**Supplementary Figure 2:**
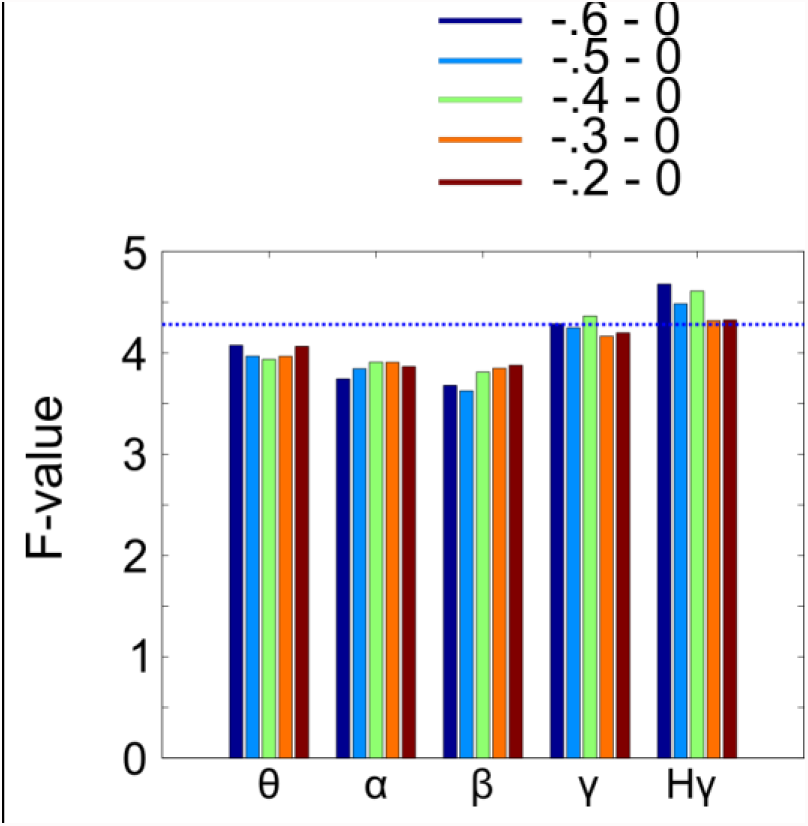
We averaged the F_interaction_ values in 5 different prestimulus intervals across the best 5% of electrodes. Only high γ activity shows significant prediction signals in each of the five prestimulus intervals as indicated by the averaged F_interaction_ values exceeding the confidence interval of the surrogate distribution derived from a permutation procedure in which trial labels (standard, deviant, regular, irregular) were randomly reassigned.

